# Image-based 3D active sample stabilization on the nanometer scale for optical microscopy

**DOI:** 10.1101/2025.01.10.629211

**Authors:** Jakob Vorlaufer, Nikolai Semenov, Caroline Kreuzinger, Manjunath G. Javoor, Bettina Zens, Nathalie Agudelo Dueñas, Mojtaba R. Tavakoli, Marek Šuplata, Wiebke Jahr, Julia Lyudchik, Andreas Wartak, Florian Schur, Johann G. Danzl

## Abstract

Super-resolution microscopy often entails long acquisition times of minutes to hours. Since drifts during the acquisition adversely affect data quality, active sample stabilization is commonly used for some of these techniques to reach their full potential. While drifts in the lateral plane can often be corrected after acquisition, this is not always possible or may come with drawbacks. Therefore, it is appealing to stabilize sample position in three dimensions during acquisition. Various schemes for active sample stabilization have been demonstrated previously, with some reaching sub-nm stability in three dimensions. However, these high-performance implementations significantly added to the complexity of the hardware and/or sample preparation. Here, we present a scheme for active drift correction that delivers the nm-scale 3D stability demanded by state-of-the-art super-resolution techniques and is straightforward to implement. Using a refined algorithm that does not depend on sparse peaks typically provided by fiducial markers added to the sample, we stabilized our sample position to ∼1 nm in 3D using objective lenses both with high and low numerical aperture. Our implementation requires only the addition of a standard widefield imaging path and we provide an open-source control software with graphical user interface to facilitate easy adoption of the module. Finally, we demonstrate how this has the potential to enhance data collection for diffraction-limited and super-resolution imaging techniques using single-molecule localization microscopy and cryo-confocal imaging as showcases.

**Why it matters:** Super-resolution light microscopy has enabled the visualization of biological structures down to the nm-scale. However, uncorrected drifts during often extended acquisition times may adversely affect data quality. Active drift correction in three dimensions has achieved sub-nm stabilization, but state-of-the-art techniques come with considerable overhead on sample preparation and/or hardware. Here, we demonstrate an image-based stabilization scheme which allows for flexibility regarding structures used for stabilization and is straightforward to adopt. Using a maximally simple implementation, we stabilized the position of our sample to around 1 nm over extended acquisition times and demonstrated usefulness in two example imaging settings where sample drifts are critical, super-resolution single-molecule localization microscopy and confocal imaging at cryogenic temperatures.

## Introduction

Fluorescence imaging is a powerful technique to reveal biological information, with super-resolution approaches (Huang et al., 2009; Sahl et al., 2017) routinely allowing for the investigation of length scales down to few 10s of nm. However, uncorrected motion of the sample during acquisition adversely affects data quality both in conventional and in super-resolution microscopy. This issue is typically aggravated for super-resolution approaches due to the higher spatial resolution and extended acquisition times.

Depending on the modality of image formation, drifts affect different super-resolution techniques in distinct ways. In single-molecule localization microscopy (SMLM) (Lelek et al., 2021) emitters are activated in a spatially stochastic manner and individually localized. Therefore, uncorrected sample movements corrupt information about the relative positions of localized fluorophores, effectively decreasing the resolution of a reconstructed dataset. Since axial drifts directly hamper the localization by defocusing the signal from single emitters, focus stabilization systems are common components of state-of-the-art SMLM setups (Lelek et al., 2021; Power et al., 2024). Compared to axial drifts, lateral drifts can be more readily corrected after acquisition, for example using redundant cross-correlation (RCC) (Wang et al., 2014) or fiducial markers (Balinovic et al., 2019; Lee et al., 2012; Li et al., 2023). However, this approach becomes more elaborate when applications require high accuracy. For example, to achieve sub-nm resolution in advanced variants of SMLM, a combination of RCC- and fiducial-based drift correction followed by local refinement was required (Reinhardt et al., 2023; Weisenburger et al., 2017). It is attractive to actively stabilize the sample in three dimensions during an acquisition, as this generates a markedly improved starting point for any downstream corrections. Moreover, active 3D stabilization allows for the observation of how the reconstructed structure builds up in real-time, avoids regions of interest to drift out of the field of view, and eliminates blurring of the point spread function (PSF) by drifts occurring within individual camera frames.

Recently, the MINFLUX (Balzarotti et al., 2017) and MINSTED (Weber et al., 2021) concepts have achieved imaging and tracking at (sub-)nanometer localization precision by combining coordinate-stochastic activation of fluorophores with coordinate-targeted readout of their spatial coordinates. These concepts crucially depend on accurately positioning light patterns with respect to fluorophore positions. Since relative drifts between fluorophore coordinates and positions of light pattern cannot be readily measured or corrected, it is essential to actively stabilize sample position in 3D during acquisition in such modalities. Under optimal imaging conditions, the precision of sample stabilization can indeed become limiting for the achievable resolution (Weber et al., 2023).

Sample motion during imaging has much broader implications, reaching beyond SMLM and nm-scale imaging. For diffraction-limited and super-resolution imaging realized on point-scanning microscopy platforms, including e.g. typical implementations of stimulated emission depletion (STED) super-resolution microscopy (Hell and Wichmann, 1994; Klar et al., 2000), drifts skew the resulting image in an unpredictable manner. Since these distortions cannot be corrected after an acquisition, drifts are problematic for accurately extracting the relationship between coordinates queried at different time points. Depending on the magnitude and timescale of drifts, they may either interfere with visualization of individual (sub-diffraction) structures or lead to larger-scale distortions that may e.g. compromise correlation to other imaging modalities, such as electron microscopy (Hauser et al., 2017).

Imaging at cryogenic temperatures offers the possibility to directly analyze structures preserved in a near-native state by rapid freezing avoiding formation of crystalline ice (“vitrification”) without the need of chemical fixation. Here, the temperature of the sample needs to be kept below ∼135 K to avoid devitrification. In this imaging setting, prominent mechanical drifts may occur due to limited mechanical stability of commonly used cryo-stages, which may impact the performance of both diffraction-limited, in particular point-scanning approaches, as well as super-resolution imaging (Dahlberg and Moerner, 2021). In addition to the development of highly stable cryo sample stages for optical imaging, there is an opportunity to increase imaging performance and correlation accuracy with potent stabilization techniques.

Active drift correction schemes capable of stabilizing sample position to (sub-)nm levels in 3D typically utilize external fiducial markers (Carter et al., 2007; Coelho et al., 2021, 2020; Grover et al., 2015; Koo et al., 2013; Schmidt et al., 2021). Adding fiducials entails an extra step in the sample preparation that might be undesirable for certain preparations or require careful optimization. Fiducial markers may show bleaching, intensity fluctuations or aggregation (Balinovic et al., 2019), as well as movements on the nm-scale relative to the structures of interest (Li et al., 2023). This prompted e.g. the development of DNA-origamis specifically tailored for nanoscale PAINT-imaging, featuring integrated binding sites for single fluorophores used as markers for drift correction (Dai et al., 2016). Fiducials in the same spectral channel(s) as the biological structures of interest may decrease usable field of view or cause out-of-focus background. Taken together, it would be desirable to have options for stabilizing either on structures directly present in the biological specimen itself or another reference structure in highly stable relation to the sample. Implementations using biological structures for stabilization have been limited by the low contrast of brightfield imaging (McGorty et al., 2013), although (Shang et al., 2021) recently achieved 3-6 nm stabilization precision using differential phase contrast imaging. Overall, currently available schemes that realize nanoscale active stabilization in three dimensions add a substantial amount of complexity to the hardware and/or sample preparation process.

Here, we describe a scheme for active stabilization of sample position capable of reaching nanoscale precision in 3D with minimal overhead to the microscope hardware and sample preparation. Our approach reaches sub-nm 3D stabilization precision with an image-based approach and is applicable also in absence of sparse signal peaks typically provided by fiducial beads. It reaches such high performance also in imaging settings using an objective lens with comparatively low numerical aperture (NA), here choosing NA 0.75. This combination of nanoscale stability with flexibility regarding sample preparation and NA promises to be an enabling feature for various techniques for which active stabilization is currently challenging. We achieved this performance by combining cross-correlation for determining lateral displacements with estimation of axial displacements through analysis of pixel intensity distributions, in both cases comparing measurement image frames to reference stacks along the respective directions.

We show the usefulness of our scheme by applying it in two example imaging settings: SMLM of the nuclear pore complex and cryo-confocal imaging of fluorescent beads. Our concept is remarkably straightforward to implement (Fig. 1A), and we provide an open-source software package including a graphical user interface to enable other researchers to adopt our method easily.

**Fig 1:**
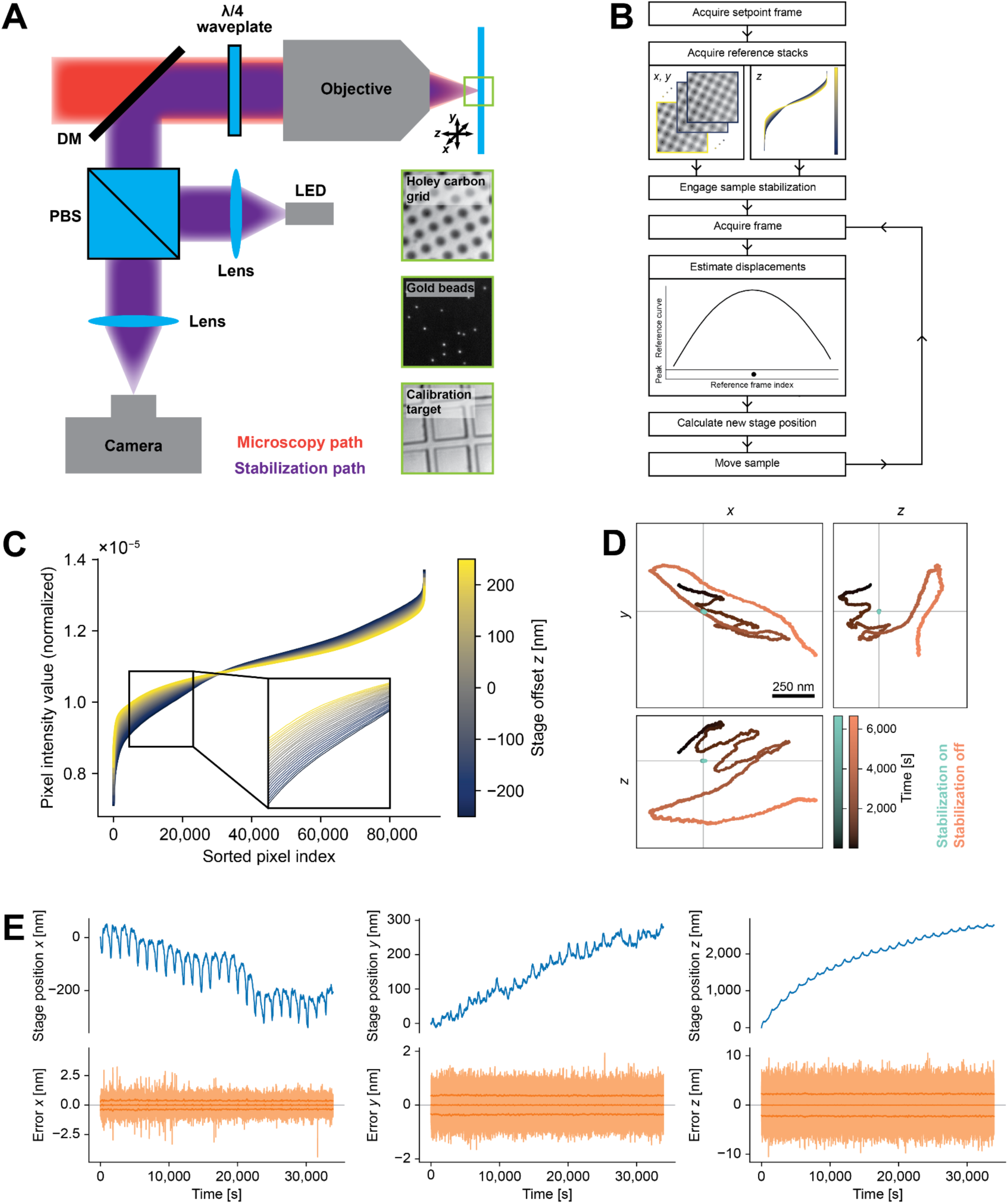
Active Stabilization Concept. **A** Schematics of optical path for sample stabilization (purple), comprising a wide-field imaging path using near infrared light, combined with the microscopy path (red) via a dichroic mirror. For further details see Fig. S1. *Right*: representative raw images of different structures used for stabilization: *Top*: holey carbon film commonly used in cryo-fluorescence imaging and EM. *Middle*: gold beads immobilized on a coverslip. *Bottom*: Magnification calibration target with 10 μm large tiles. Individual images have approx. 23 µm edge length. **B** Stabilization workflow. After choosing a ROI and acquisition of reference stacks, the scheme estimates displacements in each iteration of the active feedback loop by comparing camera frames against the reference stacks and actuating on sample position. **C** Normalized pixel intensity as a function of intensity-sorted pixel index for each frame (color-coded) of the reference stack along the z-direction. Enlarged view of the data in panel B. Curves for the individual frames of the reference stack (spaced 20 nm along *z*) can be discerned as separate lines in the magnified view. **D** Position of an individual gold bead on a coverslip over time with and without sample stabilization, tracked in the microscopy path for 1h 51 min. In the stabilization path, a distinct set of gold beads on the same coverslip was used for stabilization of sample position. **E** Long-term stabilization measurement on a holey carbon film. The blue curves show the stage movement applied to compensate for drifts. Error signals reflecting the residual deviations from the initial sample position (semi-transparent orange curves), are centered around 0. Residual deviations can be attributed mainly to noise associated with estimating displacements. The central solid orange line shows the rolling average of the error signals across 1000 data points. The lines above and below represent the standard deviation across the same window. This measurement exhibited relatively large drifts attributed to thermal equilibration (mainly along *z*) and periodic fluctuations of the laboratory temperature (mainly along *x* and *z*). Even in these suboptimal conditions, our module stabilized the sample on the nanoscale for the measurement period of more than 9 h.

## Materials and Methods

### Microscope Setup

We performed all measurements on a homebuilt setup comprising imaging paths for active sample stabilization, widefield and confocal imaging. See Fig. S1 for a schematic of the beam path as well as a list of the components used. The hardware was controlled by a standard microscope control PC equipped with an Intel Core i7 CPU (8×3.6 GHz) and 64 GB RAM.

The setup was constructed in an upright configuration to facilitate compatibility with a commercial cryo-microscopy stage (CMS196 V3, Linkam Scientific Instruments Ltd., Redhill, United Kingdom). The sample stage was mounted on a calibrated 3D piezo-stage which was used for drift correction. Light was collected by the same objective as was used for illumination. Depending on the application, we used either an oil immersion objective (100x/NA 1.45, Olympus) or a long working distance air objective (100x/NA 0.75, Leica). We used a Leica tube lens for all measurements, as this provided adequate performance also when using the Olympus oil immersion objective.

The common beam path shared by the stabilization and the other imaging modalities comprised an achromatic λ/4 waveplate placed near the objective lens to create circularly polarized light at the sample, as well as a telescope consisting of the tube lens and an achromatic doublet lens. Finally, a shortpass dichroic mirror (DM) separated the path used for sample stabilization (operated at near-infrared wavelength) from the main part of the microscope (termed “microscopy path” in Fig. 1A). At the position of the DM, the light from the sample plane was collimated to prevent aberrations in the “microscopy path” from transmission through the glass substrate. This condition could not be simultaneously fulfilled for the widefield illumination, but aberrations in this path do not directly affect image quality.

### Sample lock path

Light from a fiber-coupled LED emitting around 940 nm was collimated and spectrally filtered. We chose this spectral window because it is not commonly used in fluorescence imaging. The vertical polarization component was reflected by a polarizing beam splitter (PBS) and directed to the common optical path described above, while the horizontal polarization was dumped. The light reflected by the sample had an orthogonal polarization after passing the λ/4 waveplate near the objective lens in both directions, and was hence transmitted by the PBS. A longpass filter blocked residual light from the “microscopy path”, before the light was focused on a camera. By default, we mostly used a temperature-stabilized CCD camera at 10 ms exposure time. However, we also confirmed that the concept worked equally well, in fact with even better in-loop performance, using an industry-grade CMOS camera (Fig. 1D, S2) after allowing enough time for the camera temperature to equilibrate. Without this warm-up phase, which typically lasted for several hours depending on the heatsink attached to the camera (Fig. S2), pixel intensity values increased gradually during acquisition of reference stacks and active stabilization, which led to non-optimal performance despite intensity normalization.

The image pixel size of the employed CCD camera was 75 nm/78 nm along *x/y* directions for the air objective, and 67 nm/70 nm for the oil immersion objective, as measured with a calibration target with 10 µm tile pattern (Planotec S1934, FIAS, Biedermannsdorf, Austria). Note that objective lenses were not fully corrected for the near-infrared range. For the CMOS camera, we measured pixel sizes of 55 nm/52 nm for the air objective and 50 nm/50 nm for the oil objective (again *x*/*y*), which is in line with the difference of the specified pixel sizes of the cameras. The field of view was limited by the focusing lens in front of the camera, corresponding to a diameter of 56 μm at the sample. For the stabilization measurements, we read out a region of interest (ROI) of 300 x 300 pixels. The LED illumination delivered a maximum optical power of ∼280 μW at the sample position. We adjusted the intensity for every measurement, such that the maximum pixel intensity value was around 60-80 % of the saturation level of the camera.

### Microscopy path

For the applications tested here, we used either widefield or confocal fluorescence imaging. Motorized flip mirrors in the illumination and detection paths allowed for straightforward switching between these modalities. Laser beams at two different wavelengths (488 nm and 642 nm wavelength) were combined via DMs and coupled into one of two polarization-maintaining optical fibers for widefield and confocal illumination, depending on the position of a flip mirror.

For confocal imaging, polarization direction at the output of the fiber was chosen horizontal, light was collimated and transmitted through a PBS. The light was then reflected by a multi-band DM, and passed a tip/tilt piezo mirror which was situated in a plane conjugate to the objective’s back-focal plane in order to scan the beam across the sample. The fluorescence was collected by the same path, transmitted through the multi-pass DM and focused on a pinhole. After spectral filtering, it was detected by an avalanche photo-diode operated in single-photon counting mode. Confocal measurements were performed using home-written software for microscope control.

Light for widefield imaging exited the respective fiber with vertical polarization. It was combined with the confocal imaging path by reflecting it from the PBS that transmitted the light for confocal imaging. In the widefield illumination path, a lens before the PBS focused the light at the back focal plane of the objective lens. For widefield imaging, a flip mirror after the multi-pass DM guided the emitted light to an sCMOS camera. For evaluating the out-of-loop stability in 3D, we induced astigmatism by placing a pair of cylindrical lenses with f = ± 1 m rotated against each other before the camera, as described by Power *et al*. (Power et al., 2024). Without the added astigmatism, images had a pixel size of 53 nm for the oil objective (using 642 nm light) and 60 nm for the air objective (using 488 nm light).

### Stabilization Workflow

Our stabilization unit was controlled by a custom graphical user interface (GUI) written in Python. The GUI controlled servo motors and a piezo stage for coarse and fine adjustment of sample position, respectively, as well as the camera. It allowed for setting and saving of all stabilization parameters. Options for saving the raw image, stage position and error signals for every iteration of the feedback loop were also included. Plots showing the stage position and error signal for a given axis in real time facilitated straightforward assessment of the stabilization performance. Computer code for sample stabilization is part of this submission and is available via GitHub (see Code Availability Statement).

After defining a ROI and the stabilization parameters, the sample stabilization was engaged. First, we saved a camera frame as the setpoint of the stabilization (“setpoint frame”). Subsequently, a reference stack was acquired for every axis by moving the stage over a user-defined range (symmetrical with respect to the starting position of the stage) and step size. After every step, we allowed the stage to settle, including an additional 100 ms buffer time, and added the latest camera frame to the respective reference stack. Normalizing the frames in terms of intensity turned out to be critical for adequate performance. We achieved this by dividing each pixel intensity value by the sum across all pixels.

After the acquisition of the reference stacks, the stage was moved back to the starting position and the actual drift compensation started. The feedback mechanism iteratively estimated the amount the sample had moved since the beginning of the measurement, and actuated the sample piezo stage to compensate for these displacements. Drifts during the acquisition of reference stacks or potential systematic biases of the displacement estimation may result in constant offsets of the calculated displacements. Directly using these displacements may therefore hold the sample at a constant position that is slightly offset from the user-defined setpoint. To avoid this, our error signals were defined as the displacements with respect to the setpoint frame acquired before the reference stacks (instead of the center of the reference stacks). The algorithm used for continuous drift correction is described in the Results section.

For fine calibration of error signal amplitude in physical length units (nm), we acquired “calibration measurements” after every “stabilization measurement” shown in Fig. 2 by switching off the feedback and recording the error signal as we stepped the stage back and forth by 20 nm along the respective axis. This procedure was repeated for all axes. Due to drifts occurring during these calibration measurements, we found that it was convenient to find the calibration factors by manually tuning them. When using Gaussian fitting for displacement estimation, we typically found best correspondence between step size of the stage and error signal amplitude with scaling factors between 0.85 and 1.1. We estimate that our calibrations and hence values of the error signals stated in this publication were accurate within a few percent. The following is a pseudocode implementation of the key elements of the stabilization workflow as depicted in Fig. 1B:

**Figure 2:**
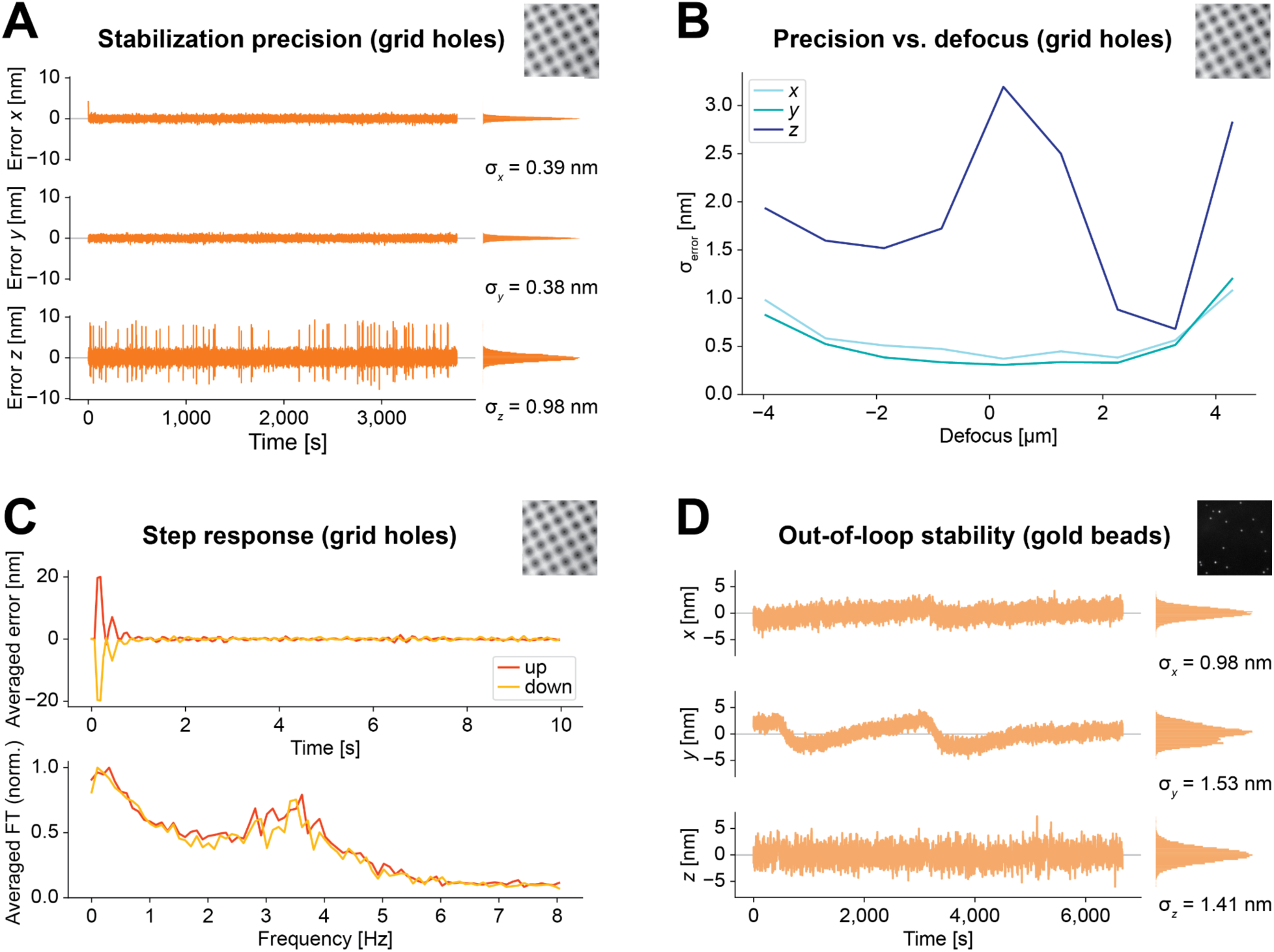
Performance Characterization. **A** Representative measurement of the in-loop stability of the active stabilization unit on holey carbon grids. The measurement was acquired at a slight positive defocus (sample shifted towards objective) with an air objective with 0.75 NA. The measured residual displacements of the sample (as given by the error signal in *x*, *y*, and *z*) showed a Gaussian distribution centered around 0 nm with standard deviation σ. Along the optical axis there were occasional spikes, which may be related to vibrations associated with the fan of the camera. **B** Precision of the active stabilization for different defocus positions using the same sample as in A. The focus position was determined by eye as the position where the structure appeared sharpest. The sample was then moved manually in steps of 1 μm along *z*, and the stabilization was engaged at each position. Each curve shows the standard deviation of the error signals (σerror) of measurements of ∼200 s duration. The lateral precision was highest in focus, and σerror remained below 1 nm within ±3 μm of the focal plane. The axial precision improved with moderate defocus. Positive defocus provided the best axial performance, but negative defocus yielded good results over a larger range. **C** Step response of the active sample stabilization. Dynamic response of the system tested by applying offsets to the stage position in steps of ±20 nm every 10 s. The error signals were reduced to the noise level within less than 1 s. The Fourier transforms (FT) show that with the settings used here, the step response produced a peak at around 3.5 Hz (0.29 s periodicity). In this particular measurement, we optimized feedback parameters for speed, while in other measurements we usually used parameters optimized for high static stabilization precision. Curves shown here represent averages of 10 steps in each direction along the *x*-axis. Raw data as well as corresponding measurements for the other axes are shown in Fig. S4. **D** Position of an individual gold bead on a coverslip over time, imaged in the microscopy path. In the stabilization path, a set of gold beads in a different region of interest was used for stabilizing sample position. The position of the bead was stabilized on the nm-scale. Residual drifts included relative drifts of the microscopy vs. stabilization path of the microscope.

#### Acquisition of reference stacks

Stage positions for reference stacks for each axis are specified by the stack range and step size. For the *z*-axis, frames are transformed into a 1D array (flattened) and pixel intensity values sorted before appending to the reference stack. For the other axes, raw images are appended to the stack.

~~~
 **for each** axis:
  **for each** specified stage position: move stage to new stage position acquire frame
    normalize frame
    **if** axis is z-axis:
    convert frame to 1D array of pixel intensity values sort array
    append sorted array to reference stack
**else:**
 append frame to reference stack
~~~

#### Displacement estimation in *x- and y-*directions

Cross-correlation CC between images *a* and *b* is calculated as follows, with the number of pixels in each row *n*_row_ and in each column *n*_col_ and the respective pixel indices *i* and *j*:

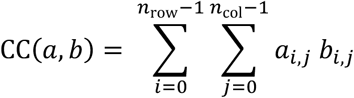

~~~
**for each** entry **in** reference stack:
  calculate CC(current frame, reference stack entry)
  append to correlation curve
scale correlation curve to range [0,1]
fit Gaussian to scaled correlation curve
displacement = center position of Gaussian
~~~

#### Displacement estimation *z-*direction

Mean squared error (MSE) between 1D arrays *a* and *b* of length *n* with indices *i* are calculated as follows:

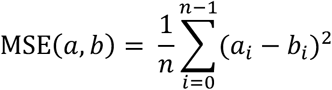

~~~
normalize frame
convert frame to 1D array of pixel intensity values sort array
**for each** entry **in** reference stack:
 calculate MSE(current array, reference stack entry) append to MSE curve
 invert MSE curve by multiplication with −1
 scale inverted MSE curve to range [0,1]
 fit Gaussian to scaled MSE curve
 displacement = center position of Gaussian
~~~

## Sample Preparation, Data Acquisition and Analysis

### Performance characterization

For the performance measurements on electron microscopy grids, we used a clipped holey carbon 2/2 200 mesh copper grid (N1-C16nCu20-01, Quantifoil Micro Tools GmbH, Großlöbichau, Germany; 2 µm hole diameter with 2 µm distances between holes) attached to a microscopy slide with parafilm. For the out-of-loop measurements on gold beads, we used a commercial alignment sample with 150 nm beads fixed on a coverslip (Abberior GmbH, Göttingen, Germany). Data analysis was mainly performed via custom Python scripts.

For out-of-loop measurements (Fig. 1D, 2C), we stabilized the sample position using a set of gold beads and additionally detected gold beads in a different region on the sCMOS camera of the widefield microscopy path with the cylindrical lens unit in place, acquiring 20,000 frames at 3 Hz frame rate with 50 ms exposure time. Here, we employed the Fit3Dcspline software package (Li et al., 2018) to localize gold beads in 3D using experimental PSFs. To generate the experimental PSF model, we used three *z*-stacks of beads acquired in different regions on the same coverslip with the cylindrical lens unit in place. Each stack spanned a range of ±700 nm at a step size of 20 nm. For display purposes, only every 5^th^ datapoint is plotted in Fig. 1D.

### SMLM of nuclear pores

For fixing gold beads on coverslips, we adapted a protocol described by Balzarotti *et al*. (Balzarotti et al., 2017). Prior to seeding the cells, #1.5H coverslips (18 mm round, 0117580, Paul Marienfeld GmbH & Co. KG, Lauda-Königshofen, Germany) were cleaned with Hellmanex III (Z805939, Merck KGaA, Darmstadt, Germany; 2 % in Milli-Q water; sonicated 2×15 min, then washed with Milli-Q water), dried with nitrogen gas and coated with 0.01 % poly-L lysine (P4707, Merck). Subsequently, they were washed 3 x 2 min in Milli-Q water, and dried with nitrogen gas before incubation with 150 nm gold beads (A11-150-CIT-DIH-1-50, Nanopartz, Loveland, Colorado, US) for 10 min which had been diluted 1:5 in Milli-Q water and sonicated for 10 min. The coverslips were washed 3 x 2 min in PBS. Finally, they were placed in new, sterile 12-well plates and sterilized with UV in a cell culture hood for 1 hour before seeding cells.

We used U-2 OS cells stably expressing a Nup96-GFP fusion protein (U-2 OS-CRISPR-NUP96-mEGFP clone no.195, 300174, CLS Cell Lines Service GmbH, Eppelheim, Germany) from Thevathasan *et al*. (Thevathasan et al., 2019) and followed their protocol for nanobody staining against GFP (Pleiner et al., 2015; Thevathasan et al., 2019). In brief, we prefixed the U-2 OS cells for 30 s in transport buffer (TRB: 20 mM HEPES pH 7.5 (H3375, Merck), 110 mM potassium acetate (4986.1, Carl Roth), 1 mM EGTA (E3889, Merck), 250 mM sucrose (84097, Merck) in Milli-Q water) supplemented with 2.5% (w/v) formaldehyde (prepared from stock F8775, Merck), then washed with TRB 2×5 min, and permeabilized with TRB supplemented with 25 μg/ml digitonin (D141, Merck) for 8 min on ice. We washed the samples 2×5 min with TBA buffer (1 % w/v bovine serum albumin (A1391, AppliChem GmbH, Darmstadt, Germany) added to TRB), and stained them with FluoTag-X4 anti-GFP nanobodies conjugated to Alexa Fluor 647 dyes (N0304-AF647-L, NanoTag Biotechnologies GmbH, Göttingen, Germany) diluted 1:250 in TBA. Cells were washed in TBA 2×5 min, then again fixed in TBA supplemented with 2.5 % formaldehyde for 10 min and washed again in TBA 2×5 min. Subsequently, 0.4 % (v/v) Triton (X100, Merck) in PBS was applied for 3 min to permeabilize the nuclear envelope. The samples were washed in PBS 2×5 min before performing another round of nanobody-staining the same way as the first time. Finally, the samples were washed 3×10 min in PBS and mounted in dSTORM buffer (500 mM TRIS (T1503, Merck), 10 mM NaCl (S7653, Merck), 10 % (w/v) glucose (G8270, Merck), 0.4 mg/ml glucose oxidase (G2133, Merck), 64 μg/ml catalase (C30, Merck) in 1x PBS) on cavity slides (1320002, Marienfeld). Immediately after mounting, the coverslips were sealed with twinsil extra-hart (Picodent, Wipperfürth, Germany).

For every dSTORM-measurement (Fig. 3), we acquired 20,000 frames at 6.7 Hz frame rate with 50 ms exposure time. Our field of view was a circle with a diameter of roughly 60 μm, limited by the aperture of the focusing lens in front of the camera. Typically, measurements contained 2-4 nuclei within the field of view.

**Figure 3:**
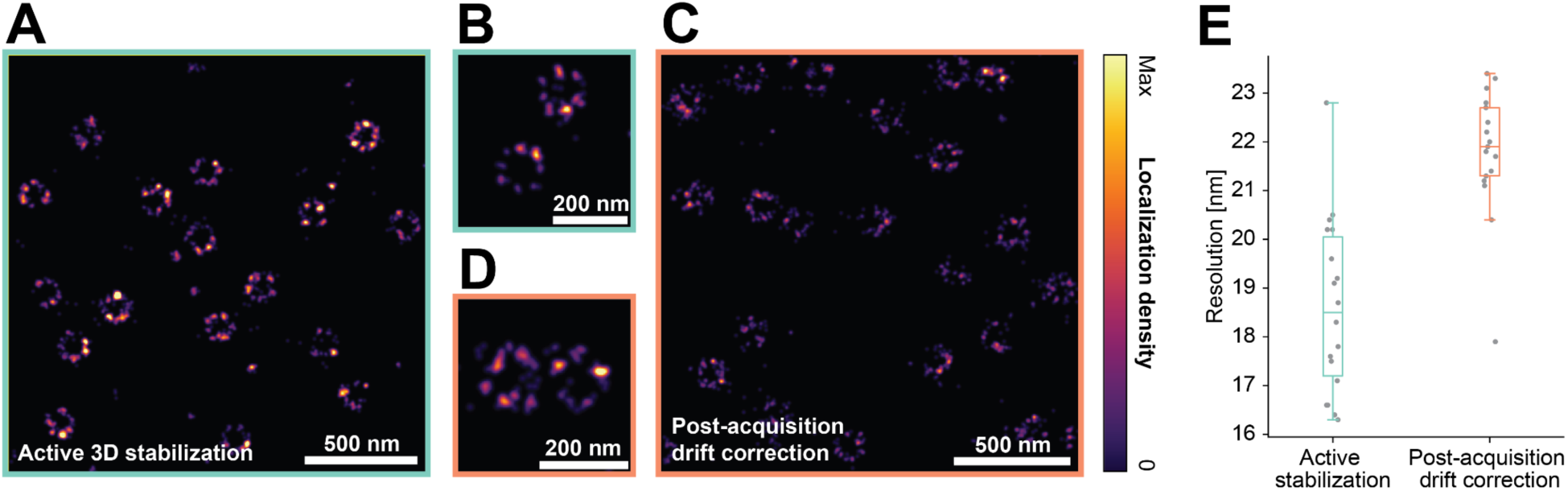
Active stabilization for dSTORM imaging of nuclear pores. **A,B** Two regions of interest at different zoom factors of a dSTORM reconstruction of nuclear pores with active stabilization engaged, showing the ring-like arrangement of the subunits. Observed variability between individual pores includes biological factors and imperfections in labeling. **C,D** Analogous measurement from different cells on the same coverslip without active stabilization, using RCC-based drift correction after acquisition. The ring-like structure of the nuclear pores was again faithfully reconstructed. However, presumably due to imperfect drift correction, the subunits were not as clearly resolved as in A and B. See Fig. S5 for additional imaging data. **E** Resolution of reconstructions with active 3D stabilization or post-acquisition drift correction displayed as box plots (lower whisker: data point within 1.5x interquartile range below first quartile, first quartile, median, third quartile, upper whisker: data point within 1.5x interquartile range above third quartile). Datapoints represent individual ROIs recorded across two individual measurements for each of the two conditions on the same coverslip. See Methods for details on resolution measurements.

We performed localization using the Picasso software package (version 0.6.8) (Schnitzbauer et al., 2017). For the two experiments for which active stabilization was disabled, we corrected drifts after acquisition in Picasso using RCC. For one of the datasets, the frame segmentation parameter was set to 200 frames and for the other we used 3 rounds of RCC with a frame segmentation of 1,000 frames.

We exported the localizations to PALMsiever (version 1.0.1) (Pengo et al., 2015), and used the FIRE plugin of the package to calculate the resolution of different ROIs in our SMLM-reconstructions. FIRE is an approach for estimating the resolution of super-resolution microscopy data based on the concept of Fourier ring correlation (Nieuwenhuizen et al., 2013).

For the resolution measurements (Fig. 3E), we selected different non-overlapping ROIs across the field of view. Every ROI spanned 4.096 x 4.096 μm^2^ at a pixel size of 1 nm. We chose the ROIs such that they contained only single-molecule blinking events, and no continuously emitting bright clusters.

### Cryo-confocal imaging

We plunge-froze 3 μl of sonicated 100 nm Yellow-Green beads (F8803, Thermo Fisher Scientific, Waltham, Massachusetts, US) diluted 1:200 in Milli-Q water on holey carbon 2/2 200 mesh copper grids (Quantifoil Micro Tools GmbH) using a GP2 device (Leica Microsystems) with 3 s blotting time. The grids were stored under liquid N_2_ temperature until imaging.

We mounted the grid on the Linkam cryo stage operated at liquid N_2_ temperature, and first acquired a low-magnification overview image using a 4x objective with a field of view spanning roughly 1.5 mm (half the grid diameter). This image was used to select areas for subsequent high-magnification light microscopy acquisition.

In order to prevent condensation of humidity from the environment on the front lens of the high-magnification air objective (working distance 4.7 mm), we heated it with a resistive foil heater attached to the side of the objective. We applied a constant current of 300 mA to the heater, which resulted in a heating power of 1.8 W applied to the objective body. The current was chosen such that the objective temperature remained approximately at room temperature throughout operation with our open cryo-stage.

We positioned the sample while observing it in widefield mode because of the higher imaging speed and larger field of view. Since grids typically exhibit bending, the focus was set to an intermediate position where the field of view contained both beads that were slightly above and below the focal plane and we opened the pinhole to detect a larger number of beads. Subsequently, we engaged the active stabilization, switched to point-scanning mode, and acquired 40 scans with approx. 34 nm and 26 nm pixel size (along *x* and *y*) and subsequently another 40 scans without stabilization.

Our confocal setup exhibited distortions due to the delayed response of our scan mirror. We corrected for such distortions using Fiji’s BigWarp plugin (Bogovic et al., 2016). Using a thin-plate spline transformation, we registered the first confocal scan via landmarks to a widefield image of the same region as largely distortion-free reference, and applied the same transformation to all confocal scans. To increase the precision of landmark placings, we placed them at the center position of the beads extracted by Fiji’s ThunderSTORM plugin (Ovesný et al., 2014). Beads which were not visible throughout the entire experiment because they drifted out of the field of view were manually excluded from the data displayed in Fig. 4A, B.

**Figure 4:**
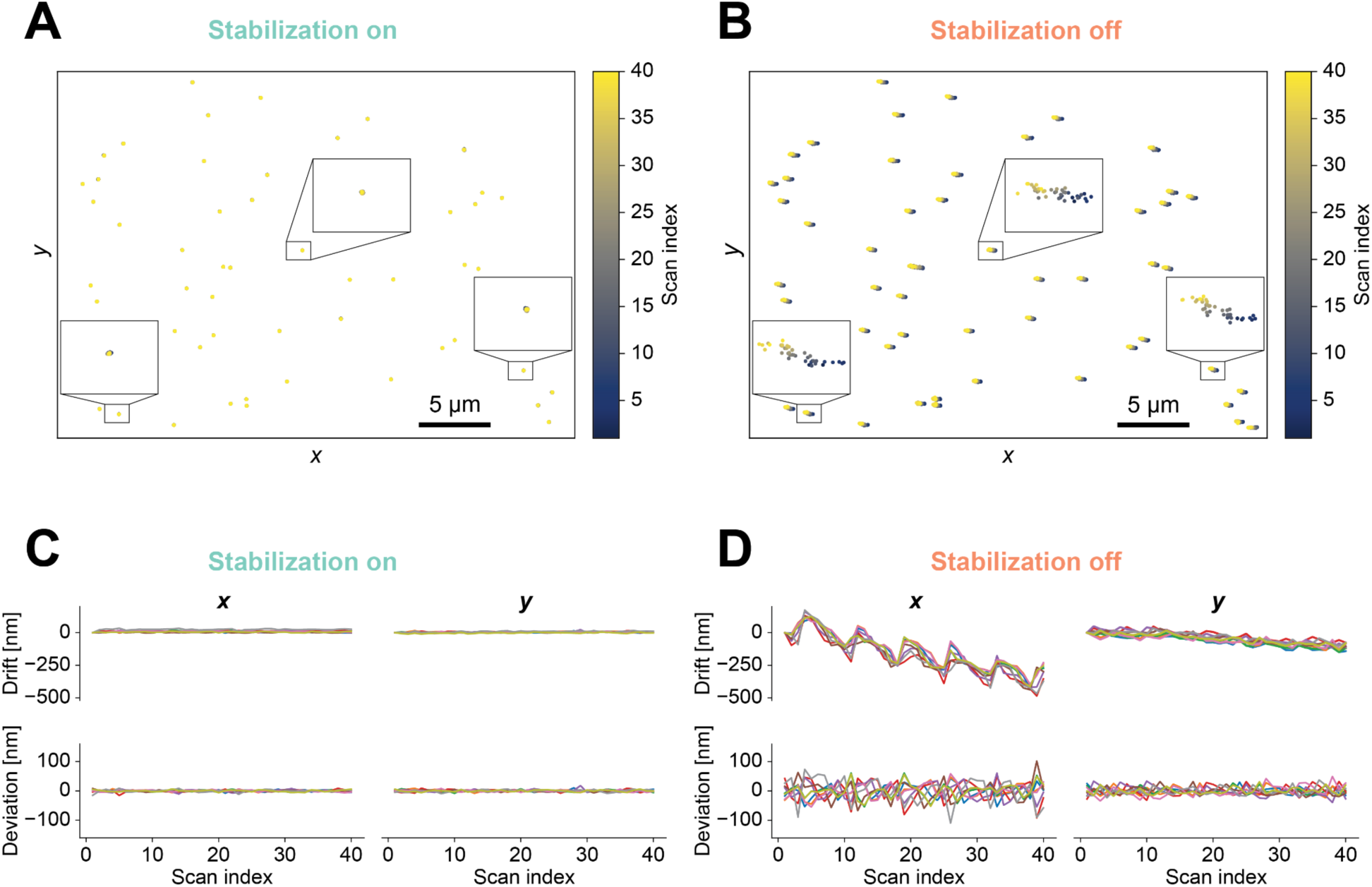
Cryo-confocal imaging. **A, B** Tracking position of fluorescent beads at cryogenic temperature over 40 confocal scans with sample stabilization engaged (A) or switched off (B), with beads embedded in vitreous ice after plunge freezing. Insets: 10-fold enlarged views of trajectories of manually selected beads. Drifts without stabilization exceeded 500 nm. Tracks differed from each other, reflecting drifts within individual scans, as beads were stably embedded in the vitreous ice. **C, D** *Top row:* Drifts of 9 manually selected beads as a function of scan index from the same datasets as A and B, respectively. *Bottom row:* Deviations of bead positions relative to the mean position of the selected beads for every scan. Display ranges of corresponding panels in **C** and **D** were identical to facilitate comparison of the associated spatial scales with and without stabilization. Active stabilization suppressed both drifts and deviations.

To show the deviation of different beads across scans (bottom row in Fig. 4C, D), we manually selected beads across the field of view. First, the localization traces were subtracted by the mean position over all scans to center them around 0. All points of the centered traces were then subtracted by the mean position of all selected beads in the respective scans to obtain deviations occurring during scans.

## Results

### Concept and Implementation

We designed our sample stabilization scheme with the aim of providing nanoscale active stabilization in all three spatial dimensions while posing minimal requirements in terms of specific hardware and modifications to the sample preparation. Sample position at the time point of engagement of the sample lock was defined as the setpoint for position stabilization, while displacement of actual position from this setpoint constituted the error signal. Error signals for stabilizing lateral (*xy*-)position were derived from image cross-correlation, whereas analysis of intensity distributions yielded the error signal for stabilizing axial (*z*-)position, both with respect to reference stacks recorded directly before engaging the stabilization. This allowed us to estimate displacements using the information encoded in all pixels within a ROI, rather than relying on peak fitting of individual fiducial markers.

We implemented our scheme by adding a standard widefield imaging path (Fig. 1A; see Methods and Fig. S1 for additional details) to a home-built microscope. To avoid interfering with fluorescence data acquisition, we chose near-infrared illumination with detection of back-scattered light on a camera. Feedback for the sample stabilization acted on a 3D piezo stage to which the sample was mounted.

The experimental workflow (Fig. 1B) started by acquiring a reference stack for every axis, which we then used for displacement estimation in the subsequent closed-loop operation. For estimating the lateral displacements at a given time, we acquired a frame and calculated the cross-correlation to all images in the reference stacks for *x*- and *y*-axes after intensity normalization. This resulted in a correlation curve for each axis whose peak position reflected the current sample position. We extracted displacements by fitting a Gaussian function. We observed that centroid calculation yielded similar precision but at times exhibited biases whose magnitude depended on the structures used for stabilization.

The same procedure worked in principle also for estimating displacements along the optical axis. However, the performance strongly depended on the measurement setup. For example, when using an air objective with NA of 0.75, the axial precision was typically around 20 nm (standard deviation), which was insufficient for high-performance super-resolution imaging. Inducing astigmatism in the detection path with a cylindrical lens improved this figure, but led to strong cross-talk to the other axes.

We therefore devised a refined approach for stabilization along the *z*-axis, avoiding the use of astigmatism: We observed that the shape of the distribution of pixel intensity values of individual frames continuously changed with *z*-position of the sample (Fig. 1C). For every iteration of the feedback loop, we thus calculated the mean squared error of the normalized, intensity-sorted pixel intensity values between the current frame and every plane in the reference stack along the *z*-axis. The corresponding curve yielded a dip whose minimum indicated the axial position. After normalization and inversion, we determined the position of the resulting peak by fitting a Gaussian function in the same manner as for the lateral directions.

For active feedback, we used a proportional-integral (PI) control algorithm, moving the piezo sample stage position. As expected, this proved to be more robust than pure proportional feedback, for example when changing samples or objectives. We also added a small portion of second order integrator, i.e. integration of the summed error, which proved advantageous in presence of strong drifts, arising e.g. from temperature fluctuations.

The active stabilization strongly suppressed drifts (Fig. 1D). When measured as mean standard deviation across 4 gold beads, while stabilizing on a different set of beads, drift reduction was by a factor of 50, 46 and 210 in *x*, *y* and *z*, with measured residual drifts comparable to the localization uncertainty. Fig. 1E shows how our active stabilization counteracted sample drifts over roughly 9 h. Fluctuations of the laboratory temperature during the measurement led to particularly strong drifts, but the residual displacement of the sample, as derived from the calibrated error signal, remained at the setpoint with nm-scale accuracy.

### Performance Characterization

We characterized the performance of our sample stabilization scheme on two diverse target structures, holey carbon films and sparse gold beads, to demonstrate the flexibility of the scheme. We used holey carbon films to evaluate the attainable stabilization precision, its dependence on defocus of the sample, and to characterize the dynamic step response. Gold beads allowed for an additional “out-of-loop” validation of stabilization performance by 3D localization of the sparse peaks in a separate imaging path.

We used the standard deviation of the error signal as a measure of the stabilization precision (“in-loop stability”). For this evaluation, we stabilized the 3D-position of an empty holey carbon grid. These are commonly used as sample carriers in electron microscopy and cryo-fluorescence imaging. We imaged the sample with an air objective (NA 0.75) with long working distance (4.7 mm), as required by our cryo-stage. At room temperature, we typically obtained in-loop stability better than 1 nm in the lateral plane, and around 1-4 nm along the optical axis (Fig. 2A-B). We stabilized the sample for a full day without observable degradation of precision.

Next, we examined the relationship between the stabilization precision and the defocus of the sample. The lateral precision did not change notably within 2 μm of the focal plane, and slightly deteriorated at larger defocus (Fig. 2B). By contrast, axial precision was highest at a defocus of 2-3 μm. The optimum defocus value and magnitude of improvement over in-focus stabilization changed for different sample types and objectives. We attribute the improved axial stability at moderate defocus to more pronounced variation of the intensity distributions with axial position (Fig. S3). This observation is not unique to our implementation, with several previous schemes for 3D active sample stabilization requiring defocus (Coelho et al., 2020; Koo et al., 2013; Lee et al., 2012; Pertsinidis et al., 2010).

To test the dynamic behavior of our implementation, we applied step-wise offsets of 20 nm in alternating directions to the stage position every 10 s and observed the step response of the sample stabilization. Fig. 2C shows the averaged error signals of 20 of these steps in either direction along the *x*-axis. The raw data as well as data for the other axes are displayed in Fig. S4. The stabilization brings the error back to the noise floor within less than a second. To quantify the temporal response, we fitted the average step responses to an exponentially decaying function with a temporal offset t_0_ and decay constant τ: *f*(t) = ae^-(t-t^_0_^)^τ. The results were similar for all axes with t_0_ ≈ 100 ms and τ ≈ 150 ms when the sample was operated at a defocus as described above. In focus, the step response along *z* was slower. The Fourier transform of the curves in Fig. 2C showed a peak at approximately 3.5 Hz followed by a monotonous decline, which represents time scales that are fast compared to typically observed microscope drifts. The sampling rate was around 16 Hz.

In practice, biological data are acquired in imaging paths that are only partially identical with the stabilization path. Accordingly, it is informative to evaluate the stabilization performance in a separate path, i.e. in an out-of-loop measurement, which also accounts for potential confounding factors, e.g. relative drifts between optical paths or crosstalk between axes in error signal generation. We performed out-of-loop measurements on gold beads immobilized on a coverslip. We used a region of interest for stabilization. In parallel, we localized a different set of beads in 3D on the camera of the widefield microscopy path, using backscattered light from a different light source.

Out-of-loop stability reached the nm-scale (Fig. 2D) over the 1h 51 min measurement time, with σ_x_ = 1.03 ± 0.09 nm, σ_y_ = 1.58 ± 0.03 nm, and σ_z_ = 1.51 ± 0.13 nm (mean ± standard deviation of five beads manually selected from different regions within a measurement). These numbers are limited by the finite precision of the out-of-loop localization. Averaging localizations over subsequent frames improved the values to σ_x_ = 0.60 ± 0.06 nm, σ_y_ = 1.43 ± 0.02 nm, and σ_z_ = 0.39 ± 0.11 nm (same beads as before, after averaging the localizations of 400 subsequent frames). However, some nanometer scale fluctuations on the time-scale of tens of minutes (Fig. 2D) remained that were not apparent in the error signals of the respective axes. We attribute them to drifts in the part of the imaging path that was distinct from the stabilization unit, likely related to temperature fluctuations in the laboratory. Along the optical axis, the standard deviation of the out-of-loop drift was about three times smaller than the in-loop error of the displacement estimation. This is presumably related to the fact that the value derived in-loop is dominated by noise in the error signal, which is averaged out by the integrator in the PI-feedback, showing that in certain scenarios drifts can be compensated on finer length scales than suggested by the in-loop precision.

### Example applications

We set out to test our scheme in two different application scenarios, SMLM at room temperature and confocal imaging at cryogenic temperatures. We selected those modalities with very different characteristics to demonstrate the versatility of the approach.

### Sample stabilization in single-molecule localization microscopy

To test our stabilization scheme in a typical high-performance imaging scenario, we chose a well-established single-molecule super-resolution modality, dSTORM (Heilemann et al., 2008), and applied it to a biological structure that is commonly used as test and reference structure in super-resolution imaging. We labeled a protein of the nuclear pore complex (NPC), which is a multi-protein assembly that regulates traffic between the nucleus and cytoplasm. NPCs comprise a central pore and a ring-like arrangement of proteins with 8-fold symmetry. The Nup96 protein is located in the ring region of NPCs and is a popular target for super-resolution imaging (Thevathasan et al., 2019). This protein is arranged in two circles with an average diameter of ∼108 nm (Wang et al., 2023).

We used a previously established cell line expressing a Nup96-mEGFP fusion protein at endogenous levels (Thevathasan et al., 2019). For dSTORM imaging, we labeled mEGFP with Alexa Fluor 647-conjugated nanobodies, thus minimizing displacement between the fluorophore and the biological target. For active 3D sample stabilization, we collected light scattered by gold beads attached to the coverslip carrying the cells.

We performed imaging of NPCs on the facet of the nuclear envelope facing the coverslip, which was well within the axial operating range of our sample stabilization scheme, and acquired dSTORM measurements with the active stabilization switched either on or off. We then compared the quality of the reconstructed NPC structures with active 3D stabilization engaged to reconstructions with conventional correction of lateral drifts after acquisition using RCC without active stabilization. We found the 8-fold symmetric structure of NPCs to be more faithfully represented with active stabilization (Fig. 3). To corroborate this observation independently of potential bias from manually selecting NPCs and average out variability of NPCs, we also determined spatial resolution in both acquisition modes with a Fourier ring correlation-based method (see Methods). The results confirmed that active stabilization indeed gave better resolution than post-acquisition correction (18.6 ± 1.8 nm vs. 21.8 ± 1.3 nm, respectively, mean ± standard deviation; p = 2.2*10^-5^ with Mann-Whitney U-Test using 18 and 17 ROIs from two measurements each on the same coverslip with active stabilization on/off).

Note that in the case where active sample stabilization was disengaged, the *z*-position was not actively stabilized either, which is different from common SMLM acquisitions. However, we gave the setup ample time to reach thermal equilibrium after turning on equipment and chose a day with high temperature stability in the laboratory, such that axial drifts were observed to be small. While the width of the detected single-molecule peaks increased slightly during the 50 minutes of acquisition (Fig. S5), this effect was small (6 % variability of the median peak width across all frames).

### Active stabilization for confocal imaging at cryogenic conditions

We chose cryo-confocal imaging as a second test case for our stabilization scheme. Imaging at cryogenic temperatures has the potential to reveal biological structures in a near-natively preserved state, but cryo-stages suitable for cryo-light microscopy often exhibit strong drifts at various timescales relevant for biological imaging, from sub-seconds to hours. Such drifts impact image quality and correlation accuracy to other imaging modalities. For example, sample drifts during an acquisition in a point-scanning approach, such as confocal imaging, may distort images in ways that cannot be corrected afterwards. Therefore, we reasoned that our 3D sample stabilization could enhance data collection in this application.

To evaluate our active sample stabilization scheme at cryogenic temperatures, we plunge-froze fluorescent beads on holey carbon grids. We used the structure of the holey carbon grid in scattering mode as reference for our 3D sample stabilization and imaged the fluorescent beads in confocal mode with our homebuilt cryo-imaging setup, using a commercial open cryo-stage to maintain the sample below the devitrification temperature. We acquired series of confocal scans with and without engaging the stabilization. When comparing the peak positions of individual beads across different scans (Fig. 4), active sample stabilization strongly mitigated both overall drifts (by a factor of 21, as evaluated from the root mean square values of the respective tracks) and apparent relative movements of different beads with respect to each other (by a factor of 6). Assuming that positions of fluorescent beads were fixed by the rigid character of the vitreous ice, relative movements of beads in the imaging data can be attributed to drifts occurring during individual confocal scans. Since our sample stabilization operated on time scales much faster than confocal scanning, it effectively fixed sample position during acquisition of individual confocal imaging frames.

## Discussion

We have developed an image-based scheme for nm-scale active sample stabilization in three dimensions with little overhead for hardware or sample preparation. We showed that our approach improved data quality on the example of two distinct imaging applications. Our algorithm does not require sparse fiducial beads to be added to the sample, and is expected to enable active sample stabilization for a wider range of scenarios than the showcase applications shown here. Our scheme is equally compatible with sparse peaks from fiducial beads, which we demonstrated using dSTORM imaging of nuclear pores. We showed that compared to post-acquisition drift correction, active stabilization resulted in an improved resolution of the single-molecule reconstructions. Moreover, we showed that active sample stabilization mitigated distortions in cryo-confocal imaging. Artifacts stemming from drifts that occur during scans cannot be easily corrected post hoc. Therefore, active stabilization can be crucial for reaching the full potential of cryogenic imaging and its correlation to other imaging modalities.

We chose to utilize the light scattered back from the sample because this configuration delivers a stable signal over extended time periods without being affected by photo-bleaching. Moreover, it does not require high illumination intensity, thus minimizing perturbations to the sample, for example when operating at cryogenic temperatures or investigating living specimens. Our stabilization unit comprised a widefield imaging path to maintain maximum simplicity of the implementation and minimize the length of the optical path, as drifts within the module itself would lead to erroneous repositioning of the sample. Still, for some applications, other contrast modalities that offer increased capacity for background suppression, such as second-harmonic imaging, Raman or fluorescence, may be preferred for generating the error signal. Also phase contrast, differential interference contrast and interferometric scattering microscopy (Kukura et al., 2009) may be applicable. Implementing our stabilization scheme with such techniques may extend the range of suitable features for stabilization to structures that produce insufficient amplitude contrast in our current configuration.

Our scheme provides a useful combination of moderate experimental complexity and modest computational requirements. Future adaptations may refine the algorithms for drift estimation to find procedures that are optimal in a mathematical sense. Alternatively, deep learning may be explored for extracting displacements. Further performance improvements may be possible if accepting additional complexity, such as use of an amplitude filter in a Fourier plane for background suppression (Schmidt et al., 2021). The microscopy path of the setup may also be actively stabilized in addition to the sample position, e.g. by using a laser beam directed at the microscopy camera as “optical fiducial” (Coelho et al., 2020; Pertsinidis et al., 2010). Overall, the design choices for a specific application will likely depend on factors such as the type of structures used for stabilization, required precision, and numerical aperture of the objective lens.

We observed that temperature stability of the camera chip was important for accurate stabilization. Therefore, we chose an actively cooled CCD camera available in the lab for our implementation. However, vibrations from the camera fan might have produced the occasional spikes visible in Fig. 2A. Using an industry-grade, passively cooled CMOS camera, we obtained even higher in-loop stabilization precision than reported in Fig. 2 with the same sample and objective (Fig. S2), after allowing for sufficient time for temperature equilibration. We reason that a camera with low readout noise and fanless active chip cooling available at moderate price would be a particularly useful choice for our scheme.

The feedback rate of our implementation depended on the acquisition parameters, including the size of the ROI and the number of frames per reference stack. By tuning these, we reached around 16 Hz update rate. At this point the readout speed of the camera became the limiting factor. While the response time was sufficient for the applications we tested, the sampling rate of the feedback can in principle be further increased by using a faster camera (suitable models can run at frame rates of 100s of Hz). Similarly, calculations for displacement estimation can be sped up by performing them on a GPU instead of a standard instrument control PC, but eventually the response time of the sample stage will limit the usable feedback rate. We did not synchronize the feedback cycles to an external clock, which, however, did not have an obvious effect on stabilization performance.

The choice of the feedback type may be affected by the desired characteristics in terms of speed and accuracy as well as the dynamics of the stabilization unit, including piezo stage, camera, PC interface, and computation. For example, while others have noted that pure proportional feedback allowed for high-performance drift correction (Coelho et al., 2020; Grover et al., 2015; McGorty et al., 2013) we opted for standard proportional/integral feedback with a small component of second-order integrator, which yielded satisfactory performance and did not require precise calibration of measured displacements.

While our implementation is based on a homebuilt setup, it should be straightforward to integrate our scheme in commercial microscopes. Various imaging modalities besides the applications shown here, for example quantitative time-lapse or multiplexed imaging, may benefit from effective, straightforward active stabilization. The ease of implementation of our scheme may reduce the entry barrier for users who may otherwise not consider active sample stabilization for their application. With its simplicity and flexibility (e.g. compatibility with high- and low-NA objective lenses), while allowing nanoscale stabilization in 3D, we expect our stabilization scheme to be of value in a range of applications.

## Supporting material

Supporting material includes five figures.

## Acknowledgements

We acknowledge expert support by ISTA’s scientific service units, including the Miba Machine Shop, the Electron Microscopy Facility, Lab Support Facility.

We gratefully acknowledge funding by the following sources.

Chan-Zuckerberg Initiative, Visual Proteomics Imaging grant 2021-23474 (FS and JGD). Austrian Science Fund (FWF) grant DK W1232 (MRT and JGD).

Austrian Academy of Sciences DOC fellowship 26137 (MRT).

Marie Skłodowska-Curie Actions Fellowship GA no. 665385 under the EU Horizon 2020 program (JL).

ISTA postdoctoral fellowship IST fellow (AW).

Human Frontier Science Program postdoctoral fellowship LT000557/2018 (WJ).

## Author Contributions

JV and JGD designed the study, interpreted data, and wrote the manuscript. JV performed experiments and analysis. NS programmed the sample stabilization user interface. CK, NAD, MGJ, BZ, MRT performed sample preparation. MŠ programmed hardware control. WJ programmed control software for spatial light modulator. AW supported hardware design. JL advised on computational implementation. FS advised on cryo-sample preparation and imaging.

## Code Availability

Computer code for sample stabilization is available at https://github.com/danzllab/samplestabilization under the GNU Affero General Public License (GNU AGPLv3) license. Copyright: Institute of Science and Technology Austria. Computer code for spatial light modulator (SLM) control is available at https://github.com/danzllab/SLMcontrol under the GNU Affero General Public License (GNU AGPLv3) license. Copyright: Institute of Science and Technology Austria.

