## Supplementary Information for "Image-based 3D active sample stabilization on the nanometer scale for optical microscopy"

### Supplementary Figure 1: Optical setup

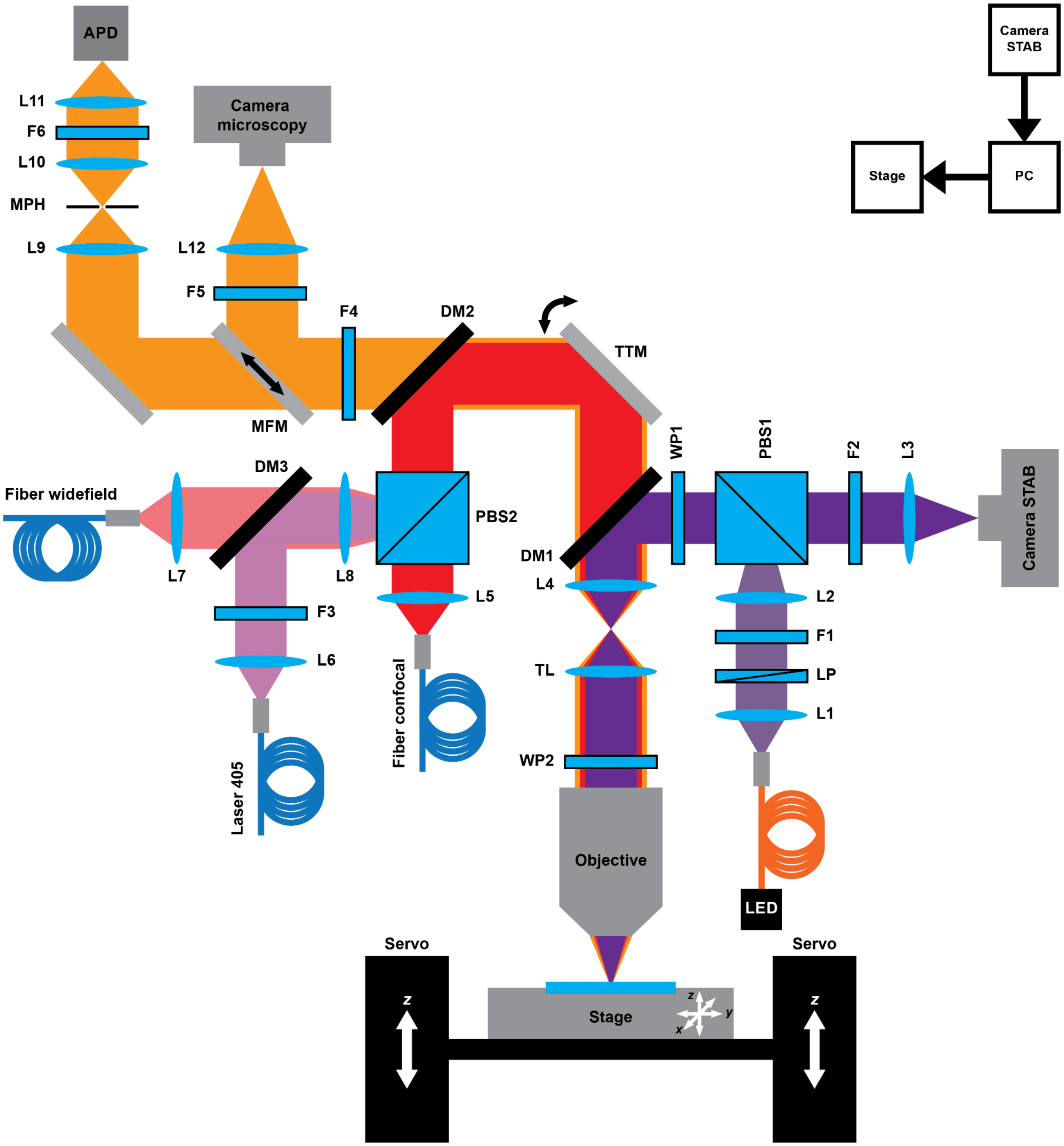


Optical path with light for sample stabilization (purple), illumination for microscopy data acquisition (red) and light scattered or emitted from the sample (orange). In the widefield and sample stabilization paths, illumination light is focused on the back focal plane of the objective and approximately collimated at the sample.

For widefield and confocal illumination, lasers were spectrally filtered, combined on a dichroic mirror and coupled into the respective polarization-maintaining single-mode fiber. For simplicity, the laser module is not depicted in the figure.

**Components:**

*Stabilization unit:*
LED: 940 nm fiber-coupled LED (M940F3, Thorlabs, Newton, New Jersey, USA) driven by LEDD1B (Thorlabs), coupled into multimode fiber with 0.22 NA (M47L01, Thorlabs).
Camera STAB: stabilization camera, CCD (Luca R, Andor, Belfast, UK) or CMOS (U3-3060SE-M-GL Rev.1.2, IDS Imaging Development Systems, Obersulm, Germany).
LP: linear polarizer (LPNIRE100-B, Thorlabs) on rotational mount.
PBS1: polarizing beamsplitter, 700-1100 nm coating (PBS252, Thorlabs).
WP1: zero-order λ/4 waveplate (WPQ10M-915, Thorlabs).
L1: achromatic doublet, f = 50 mm (AC254-050-B-ML, Thorlabs).
L2: achromatic doublet, f = 150 mm (AC254-150-B-ML, Thorlabs).
L3: achromatic doublet, f = 80 mm (AC254-080-B-ML, Thorlabs).
F1: 950/57 nm bandpass filter (ET950/57x, Chroma Technology, Bellows Falls, Vermont, USA).
F2: 808 nm longpass filter (BLP01-808R-25, Semrock, Rochester, New York, USA).

*Illumination:*
Laser 405: 405 nm activation laser, 100 mW (IBEAM-SMART-405-S, Toptica Photonics, Gräfelfing, Germany).
Fiber widefield: polarization-maintaining single-mode fiber for widefield excitation (PMC-E-400RGB-2.8-NA011-3-APC.EC/OPC.EC-400-P, Schäfter + Kirchhoff, Hamburg, Germany).
Fiber confocal: polarization-maintaining single-mode fiber for confocal excitation (QPMJ-3AF3S-405/650-3/125-3AS-5-1-WK, Oz Optics, Ottawa, Canada).
PBS2: polarizing beamsplitter, 400-700 nm coating (PTW 0.25, Bernhard Halle Nachfl., Berlin, Germany).
L5: achromatic doublet, f = 40 mm (AC254-040-A-ML, Thorlabs). Telescopes of beam modulation unit (not shown, see below) between L5 and PBS2 decrease beam diameter by 0.44x.
L6: achromatic doublet, f = 25 mm (AC127-025-A-ML, Thorlabs).
L7: achromatic doublet, f = 100 mm (AC254-100-A-ML, Thorlabs).
L8: achromatic doublet, f = 150 mm (AC254-150-A-ML, Thorlabs).
F3: 405/10 nm bandpass filter (FF01-405/10-25, Semrock).
DM2: 405/488/561/640 multipass dichroic mirror (ZT405/488/561/640rpcv2-UF3, Chroma).
DM3: 405 nm longpass dichroic mirror (F48-403, AHF Analysentechnik, Thübingen, Germany).

*Detection:*
Camera microscopy: sCMOS camera for widefield detection (Orca Fusion BT, Hamamatsu Photonics, Hamamatsu, Japan).
APD: single-photon counting module (COUNT-100B, Laser Components, Munich, Germany).
MFM: motorized flip mirror (MFF101/M, Thorlabs).
MPH: motorized pinhole (MPH16, Thorlabs).
L9: achromatic doublet, f = 100 mm (AC254-100-A-ML, Thorlabs).
L10: achromatic doublet, f = 75 mm (AC254-075-A-ML, Thorlabs).
L11: achromatic doublet, f = 30 mm (AC254-030-A-ML, Thorlabs).
L12: achromatic doublet, f = 80 mm (AC254-080-A-ML, Thorlabs); for 3D localization additionally two cylindrical lenses with f = 1000 mm (LJ1516RM-A, Thorlabs) and f = -1000 mm (LK1002RM-A, Thorlabs) on rotational mounts.
F4: 842 nm shortpass filter (FF01-842/SP-25, Semrock).
F5: 698/70 nm (FF01-698/70-25, Semrock) and 706/95 nm (ET706/95m, Chroma) bandpass filters.
F6:525/50 nm bandpass filter (FF03-525/50-25, Semrock).

*Common part:*
Objective: air objective (HC PL FLUOTAR L 100x/0.75, Leica Microsystems, Wetzlar, Germany) or oil objective (UPLXAPO100XO, Olympus, Tokyo, Japan).
DM1: 770 nm shortpass dichroic mirror (T770spxr-1380-UF3, Chroma).
WP2: achromatic λ/4 waveplate (AQWP10M-580, Thorlabs).
TL: tube lens (11090148038000, Leica).
L4: achromatic doublet, f = 75 mm (AC254-75-AB-ML, Thorlabs).
Stage: 3D sample piezo stage (P-733.3DD, Physik Instrumente (PI) GmbH & Co. KG, Karlsruhe, Germany), driven by E-727 (Physik Instrumente).
Servo: two translation stages (DRV250, Thorlabs) driven by BSC202 (Thorlabs); connected by 12.7 mm thick breadboard.

*Data acquisition:*
PC: Microscope control PC (C9Z390-CG, Super Micro Computer., San Jose, California, USA) with input/output modules for hardware control (field-programmable gate array USB-7856R, National Instruments, Austin, Texas, USA and acquisition module USB-3114, Meilhaus, Alling, Germany).

*Laser module (not shown):*
Laser 642: 642 nm excitation laser, 2W (2RU-VFL-P-2000-642-B1R, MPB Communications, Pointe-Claire, Canada).
F6: 640/10 nm bandpass filter (ZET640/10x, Chroma).
AOM: acousto-optical modulator (MT110-A1,5-VIS, AA Opto-Electronic, Orsay, France).
Laser 488: 488 nm excitation laser, 200 mW (IBEAM-SMART-488-S-HP, Toptica Photonics, Gräfelfing, Germany).
F7: 488/10 nm bandpass filter (ZET488/10x, Chroma).

*Beam modulation unit (not shown, used here to position Gaussian beam):*
The microscope setup contained further functionality for beam steering (electro-optic deflectors, M-311A, ConOptics Inc., Danbury, Connecticut, USA) which was not utilized for the measurement shown in this manuscript and beam shaping (spatial light modulator, X13268-01, Hamamatsu) used here to align the utilized Gaussian beam for confocal imaging with an adjustable blazed grating in the confocal path. The modulation unit also comprised achromatic waveplates for polarization control as well as telescopes resulting in an overall magnification of 0.44x between the collimation lens L5 and PBS2. Computer code for spatial light modulator (SLM) control is available at https://github.com/danzllab/SLMcontrol (see Code Availability Statement).

### Supplementary Figure 2: Stabilization with industry-grade CMOS camera

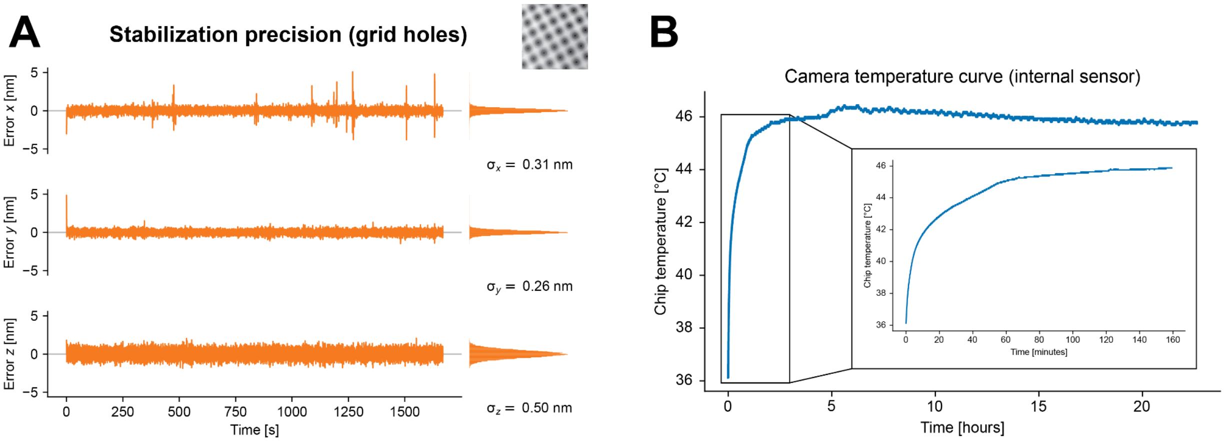


**A** In-loop stabilization measurement on holey carbon grid, similar to Fig. 2A but using an industry-grade CMOS camera without temperature stabilization. After allowing for sufficient time for the temperature of the camera chip to equilibrate, this camera reached better stabilization precision than the temperature-stabilized CCD camera used in Fig. 2A. **B** Camera chip temperature as a function of time upon start of camera acquisition. The temperature rose steeply for the first hour and reached equilibrium after ~3 hours (see magnified view of boxed region in inset). After this point, the temperature remained within less than 1 °C for 20 hours. The characteristics of this curve depend on the camera model as well as the heatsink the camera is mounted on.

### Supplementary Figure 3: Raw curves for displacement estimation

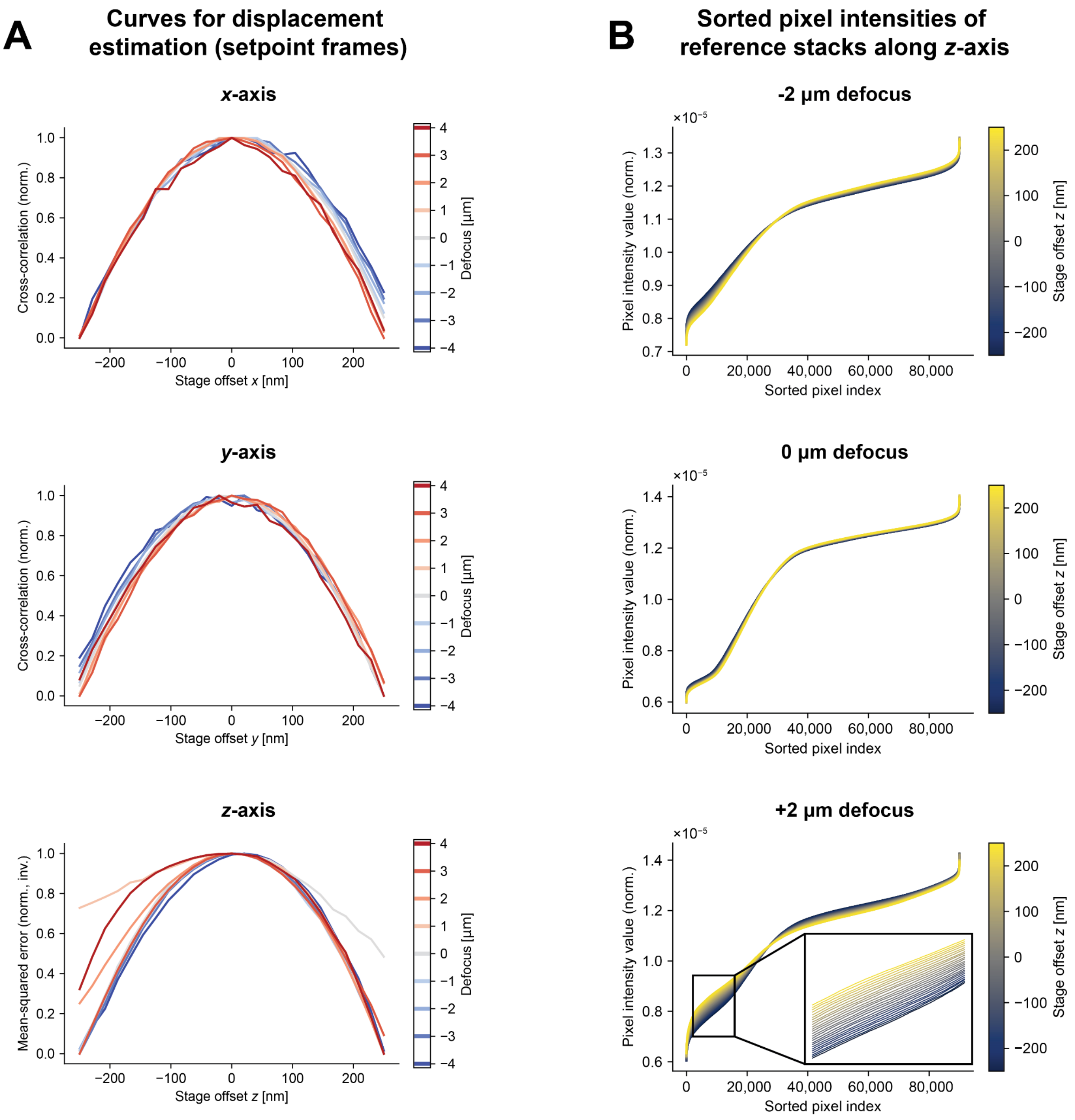


**A** Curves used for estimation of displacements on the example of the setpoint frame for the dataset in Fig. 2B. *Top, Middle*: Cross correlation between the setpoint frame and each image of the reference stack in *x*-direction and *y*-direction for different values of defocus. *Bottom*: Mean squared error (inverted) of the sorted pixel intensity values of the setpoint frame and of the individual frames of the respective reference stack for different values of defocus. We attribute slight offsets of individual peaks from the center position to drifts occurring during the acquisition of the reference stacks. In the lateral plane, the width of the peaks remained roughly constant for different values of defocus. Along the optical axis, curves exhibited different widths with varying defocus, with greater width potentially contributing to decreased precision of displacement estimations. **B** Sorted pixel intensity values for reference stacks along the *z*-axis at different values of defocus (same experiment as shown in A). The lines are color-coded according to the stage *z*-position. The visual impression shows that there is a greater variance in the defocused stacks. This is in line with the observation in A that at 0 defocus, the mean-squared error increased more slowly away from the setpoint compared to ±2 µm defocus, which led to broader peaks. Values are normalized to the sum of all pixel intensity values in the individual frames.

### Supplementary Figure 4: Step response additional data

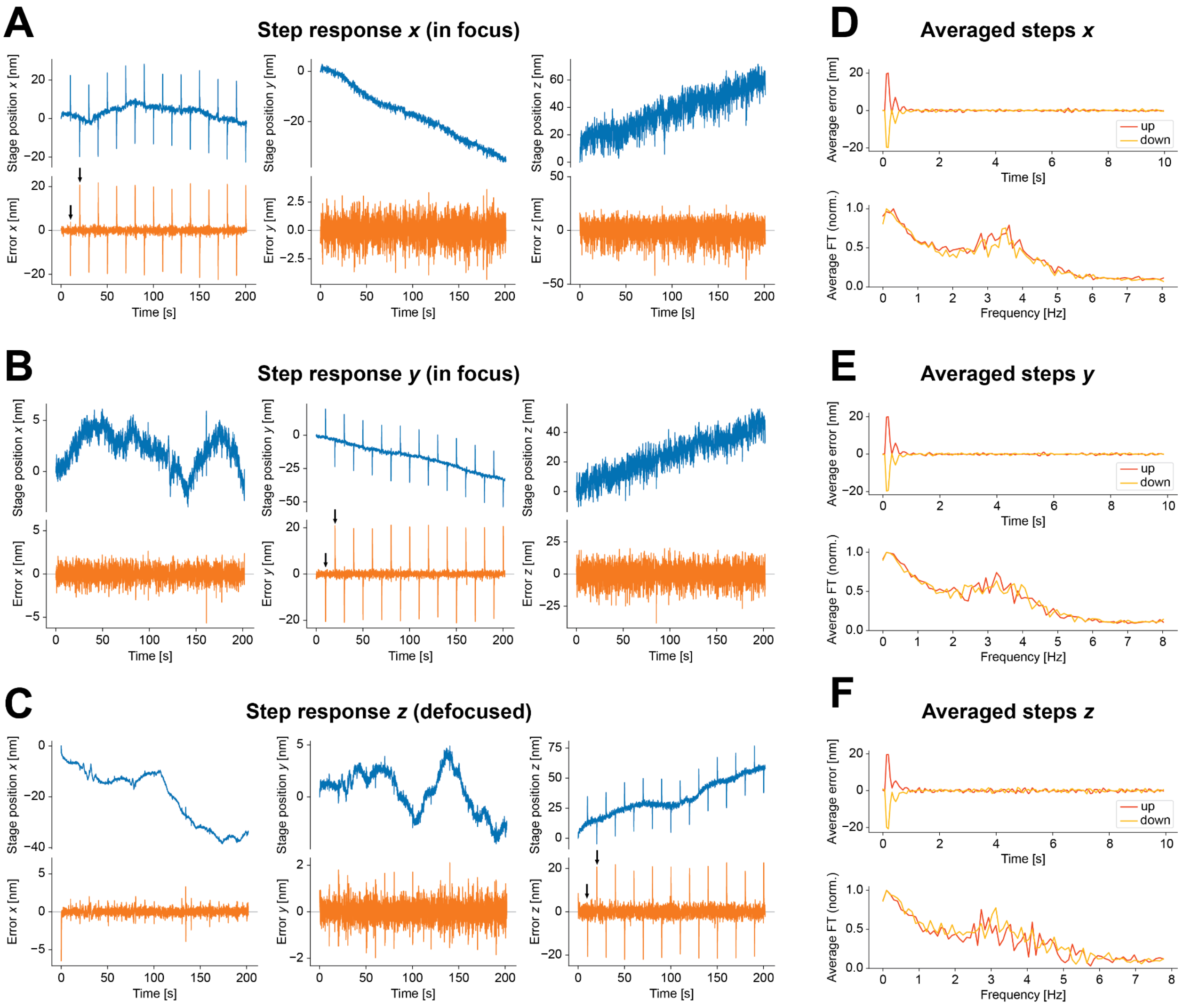


**A**, **B**, **C** Stage position (blue) and error signal (orange) during step response of the stabilization to ± 20 nm excursions applied as offsets to the stage position. Raw data for the measurements in Fig. 2C (excursion along *x*-axis) and for excursions along *y*- and *z*-axes. For lateral excursions, the measurement was performed with zero defocus, for axial excursions, a slight defocus was chosen to increase precision. The first steps in both directions are marked by black arrows. The steps produced pronounced spikes in the error signals (orange lines) along the respective axes to which they are applied. The original position was reached within less than a second. The other axes did not indicate cross-talk between the axes higher than the noise floor of the stabilization. **D**, **E**, **F** Averaged step response in time and frequency domain for steps along the *x*-direction (same data as in Fig. 2C) and *y*- and *z*-directions for the data displayed in panels A-C. Averages of 10 steps in each direction.

#### Supplementary Figure 5: dSTORM imaging of nuclear pores

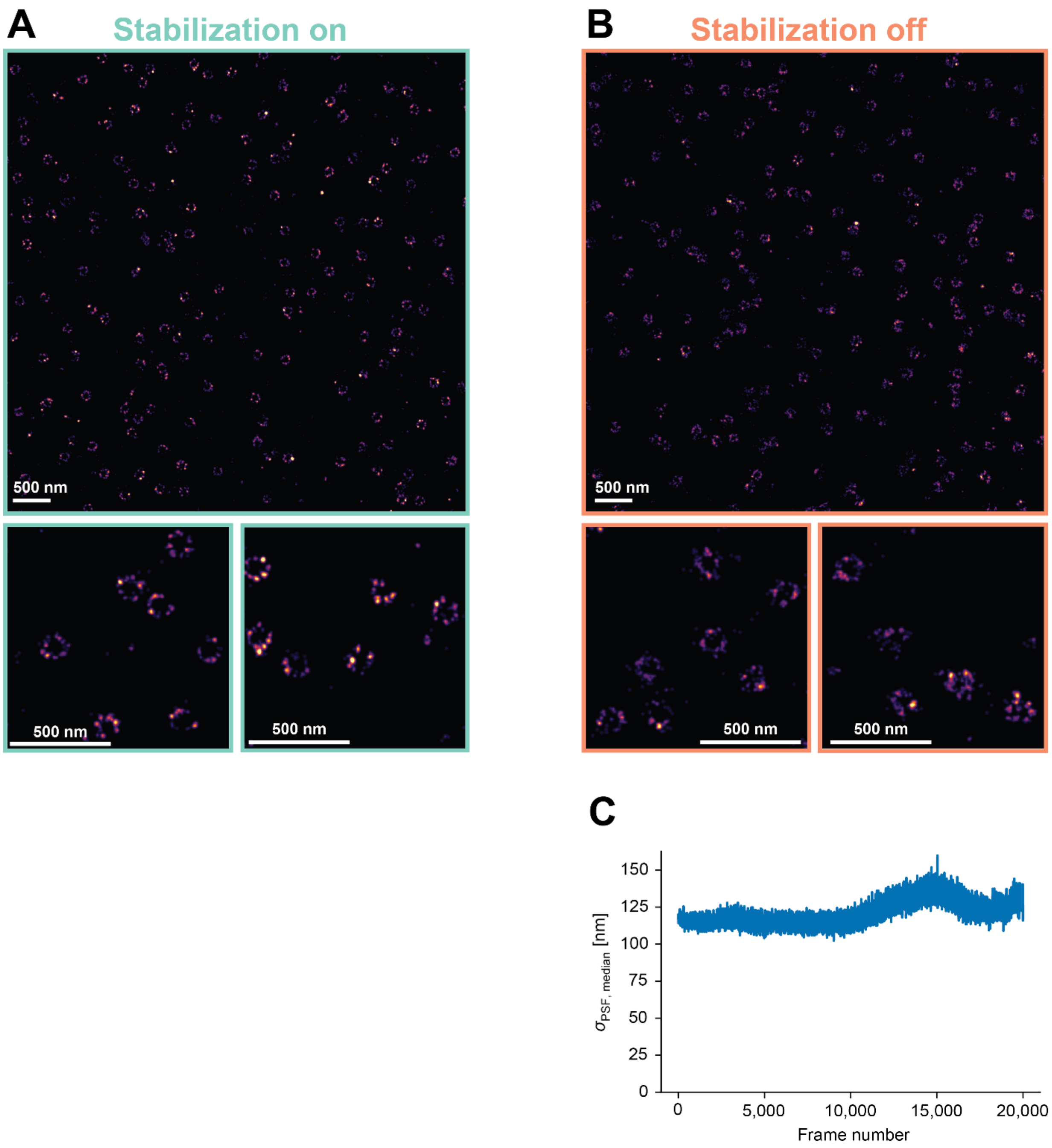


**A**, **B** Additional data comparing dSTORM acquired with 3D active sample stabilization (A) to data acquired without active stabilization but drift correction after the acquisition (B). While the ring-like arrangements of Nup96 are reliably reconstructed in both scenarios, the subunits appear crisper when active stabilization was used. Different regions of the same measurement as in Fig. 3. Data representative of two replicates. Same color map as in Fig. 3. **C** Median PSF width as a function of frame number for the measurement without active stabilization of sample position, extracted as the standard deviation $\sigma_{PSF}$ of Gaussian fits in the localization software Picasso. There was a slight widening of PSFs over the course of the measurement with a variability of ~6% (measured as standard deviation divided by the mean of $\sigma_{PSF,\ median}$ across all frames). This sets an upper bound to PSF blurring by uncorrected drifts along the optical axis and corresponding decline in localization precision.
